# Turing patterns are common but not robust

**DOI:** 10.1101/352302

**Authors:** Natalie S. Scholes, David Schnoerr, Mark Isalan, Michael Stumpf

## Abstract

Turing patterns (TPs) underlie many fundamental developmental processes, but they operate over narrow parameter ranges, raising the conundrum of how evolution can ever discover them. Here we explore TP design space to address this question and to distill design rules. We exhaustively analyze 2- and 3-node biological candidate Turing systems: crucially, network structure alone neither determines nor guarantees emergent TPs. A surprisingly large fraction (>60%) of network design space can produce TPs, but these are sensitive to even subtle changes in parameters, network structure and regulatory mechanisms. This implies that TP networks are more common than previously thought, and evolution might regularly encounter prototypic solutions. Importantly, we deduce compositional rules for TP systems that are almost necessary and sufficient (96% of TP networks contain them, and 95% of networks implementing them produce TPs). This comprehensive network atlas provides the blueprints for identifying natural TPs, and for engineering synthetic systems.

## Introduction

Pattern formation is an essential aspect of development in biology and we have a wealth of examples how complex, structured, multi-cellular organisms develop from single fertilized cells. Many organisms develop complex spatial features with exquisite precision and robustness, and this has been the subject of extensive molecular and theoretical study (Green and Sharpe, 2015; Maini et al., 2012).

Various mechanisms have been proposed to explain developmental patterning processes, ranging from maternally inherited cues (Wolpert, 1969), to mechanical forces (Howard, Grill, and Bois, 2011) and chemical reaction-diffusion networks or Turing patterns (TPs) (Gierer and Meinhardt, 1972; Turing, 1952). The latter were first proposed by Alan Turing in 1952 (Turing, 1952), and were later independently described by Gierer and Meinhardt (Gierer and Meinhardt, 1972). TPs are particularly intriguing because they are capable of generating entirely self-organized, complex, repetitive patterns of gene expression (Figure 1A, B).

TPs generally alter local concentrations of biochemical components, resulting in self-organized spatial patterns such as spots, stripes and labyrinths (Kondo and Miura, 2010). These patterns have unique and useful biological properties: perturbing them results in recovery and re-organization of the patterns (“healing”), as an intrinsic property of the dynamical biochemical interactions. This also implies that if there is variability in size across individuals, the TPs will automatically re-scale themselves, simply adding or subtracting pattern segments in response to different field sizes. This is a valuable property to support changes in size, both within existing populations and over evolutionary time. In addition, TP networks are extremely parsimonious, often employing just two or three biochemical species. This implies that they might be an economical solution for evolution to employ, wherever repetitive self-organizing patterns are needed.

Given these advantages, it is perhaps not surprising that TPs are regarded as the driving morphogenetic patterning mechanisms in many biological systems. These include bone and tooth formation, hair follicle distribution and the patterns on the skins of animals, such as fish and zebras (Raspopovic et al., 2014; Sick et al., 2006; Jung et al., 1998; Nakamasu et al., 2009; Economou et al., 2012). However, despite several experimentally verified examples (Raspopovic et al., 2014; Sick et al., 2006), the underlying complexity in biological systems has often prevented identification of the precise molecular mechanisms governing the potentially large number of TPs in nature. A second key problem is that there is a paradox between the apparent widespread distribution of natural TPs and the observation — from mathematical analyses (Gaffney, Yi, and Lee, 2016; Iron, Wei, and Winter, 2004; Palmer, 2004; Meinhardt and Gierer, 2000; Gierer and Meinhardt, 1972) — that kinetic parameters need to be finely tuned for TPs to arise. This raises the questions of how evolution could ever discover such tiny islands in parameter space and, even if it could, how would the resulting developmental mechanisms still occur robustly under noisy real conditions.

One approach to resolve these apparent contradictions is to explore TP systems mathematically. While a rich mathematical literature on Turing patterns exists, the vast majority of studies analyze single, idealized networks with fixed parameters (Gaffney, Yi, and Lee, 2016; Iron, Wei, and Winter, 2004; Liu et al., 2013). Although these studies have significantly increased our understanding of patterning mechanisms, they do not provide general guidelines for either the identification of naturally evolved Turing networks in biological systems, or for synthetic engineering of TP networks (Scholes and Isalan, 2017; Borek, Hasty, and Tsimring, 2016; Carvalho et al., 2014; Duran-Nebreda and Solé, 2016; Boehm, Grant, and Haseloff, 2018; Cachat et al., 2016; Diambra et al., 2015; Cachat et al., 2016). A recent approach which has proved very successful in increasing our understanding of biological design principles is the “network atlas” approach (Babtie, Kirk, and Stumpf, 2014; Ma et al., 2009). In this approach, biological networks that execute a particular function are modeled exhaustively: for example, all 2 and 3-node inducible genetic networks that achieve switch-like behavior were modeled by MA et al. (Ma et al., 2009). Similarly, all 3-node networks that form a central stripe pattern in a developmental morphogen signaling gradient were modeled by the group of Sharpe (Cotterell and Sharpe, 2010; Schaerli et al., 2014). Such approaches allow one to compare network and parameter design space (Barnes et al., 2011) with the resulting phenotypic map, resulting in an atlas or guidebook for the design principles behind that function. A guidebook of potential mechanisms and design rules for discovering TP networks would help towards solving the problem of characterizing molecular players in natural TP systems. Furthermore, a comprehensive Atlas of Turing network space might shed new light on the problem of how evolution could ever discover and stabilize systems which only ever function in tiny islands of parameter space.

In terms of progress towards making a TP network atlas, two recent studies have begun to analyze larger sets of TP network topologies and parameters and have made important progress in our global understanding of Turing systems (Zheng, Shao, and Ouyang, 2016; Marcon et al., 2016). Zheng et al. analyze all 2- and 3-node networks for one particular choice of regulatory function (Zheng, Shao, and Ouyang, 2016). This study does thus not consider different regulatory mechanisms, and the number of identified Turing networks is only a fraction of those identified here which is easily explained by the differences in how exhaustively the parameter-spaces are explored. In a further study Marcon et al. use exponential and sigmoidal regulatory functions and do neither include a basal production rate nor a degradation term (Marcon et al., 2016). Moreover, they shift the regulatory functions such that the stable steady states are always at zero concentrations. This allows for analytical computations and allows the authors to analyze all 3-node and 4-node networks. However, these mathematical simplifications lead to small number of networks being stable, and only a tiny fraction exhibiting Turing patterns, even compared to (Zheng, Shao, and Ouyang, 2016). But shifting the stationary states to zero has been shown to systematically misrepresent the stability properties of real dynamical systems (Kirk et al., 2015; Maclean, Kirk, and Stumpf, 2015): as this procedure destroys the dependencies between stability of stationary states and the reaction rate parameters (Kirk et al., 2015) which results in high rates (frequently up to 90% or more) of misclassifying stable stationary states as unstable.

Therefore, an exhaustive analysis and comparison of different, biologically relevant regulatory functions, as well as comprehensive sensitivity/robustness analyses of Turing networks with respect to parameter variations, still remain difficult to achieve. This is because there is a combinatorial explosion in the number of conditions to explore, and this is computationally nearly prohibitively expensive.

In this study, we develop a new computational pipeline capable of analyzing stable solutions for a wide range of potential TP network structures. Briefly, we use linear stability analysis to determine the emergence of stable TPs (see Methods) and the corresponding wavelength of the pattern (Figure 1C).

**Figure 1:**
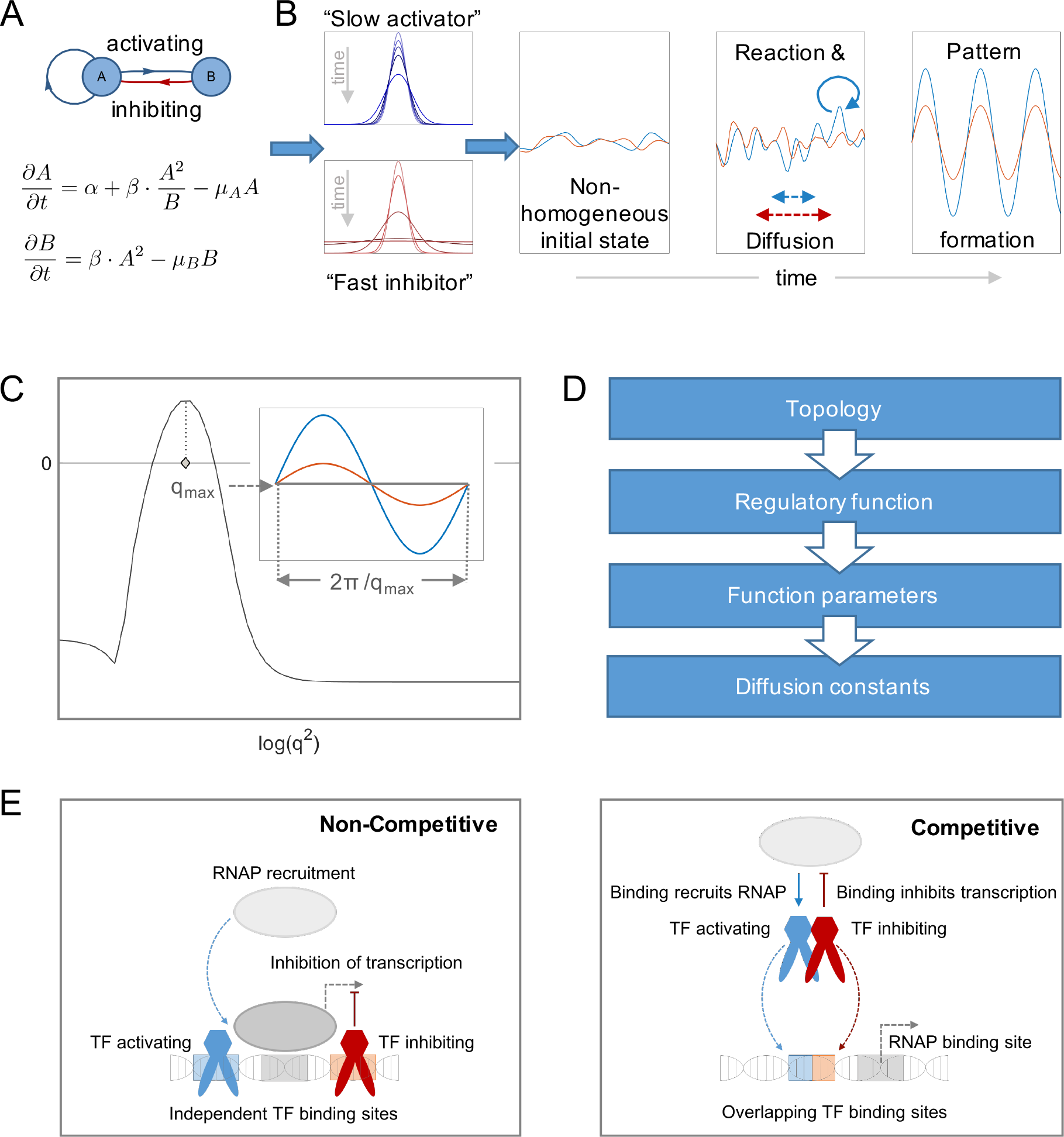
Reaction networks and Turing instabilities. (A) A network graph of the Gierer-Meinhardt model (Gierer and Meinhardt, 1972) as an example for a 2-node Turing network (top), with the corresponding ordinary differential equations below (bottom). Blue and red arrows indicate activating and inhibiting regulations, respectively. Species A activates both itself and species B, while species B inhibits species A. (B) The top left panel represents the diffusion profiles for species A (blue, slow activator) and the bottom panel for species B (red, fast inhibitor). Over time, small deviations in a noisy, non-homogeneous initial condition (Panel 2) can get amplified by the interplay of reactions and diffusion (Panel 3). For the given system this can lead to the formation of stable patterns (Panel 4). (C) An exemplary dispersion relation (real part of the largest eigenvalue of the linearized system as a function of wavenumber *q*.) of the system shown in (A). The wavenumber *q*_*max*_ for which the dispersion relation is maximal becomes amplified the strongest. This leads to the formation of a pattern with wavelength 2*π*/*q*_*max*_ as shown in the inset. (D) In this article we analyze four hierarchical factors determining a network’s pattern forming properties: the topology — the species and types of interactions between them; the regulatory function — the functional form of the interactions; kinetic parameters — parameters in the regulatory functions; and the diffusion constants of the different species. (E) Visualization of the two regulatory mechanisms analyzed in this study on a transcriptional level. Non-competitive regulation describes the case where transcription factors (TF) bind to independent TF sites and thus regulate the recruitment of RNA polymerase (RNAP) and transcription independently (left panel). In the competitive case, in contrast, TFs directly compete for the binding site.

Thus, we generate an extensive analysis of 7757 unique networks with up to three reacting species, testing them exhaustively for their ability to form TPs (we analyzed approximately 3 × 10^11^ different scenarios - amounting to 8 CPU years computing time). As summarized in Figure 1D, these are analyzed in terms of (1) the network topology; (2) the regulatory function; (3) the kinetic parameters; (4) the diffusion constants of the different species. We consider both competitive and non-competitive regulatory mechanisms (Figure 1E) and study their quantitative and qualitative differences.

In this way, we are able to systematically explore what proportion of network topologies are capable of generating TPs. Moreover, we rank these networks with respect to their robustness to variations in network topology, kinetic parameters and diffusion rates, allowing us to determine which kinds of networks are more robust. Using an unsupervised classifier, we thus identify a set of irreducible or minimal networks from which all Turing networks can systematically be constructed. This in turn allows us to distill and test compositional rules for predicting whether a given network will support TPs.

With these results in hand we can identify the networks that are most suitable for downstream synthetic engineering under different physiological conditions. There is growing interest in synthetic biology to engineer patterning systems from first principles (Basu et al., 2005; Schaerli et al., 2014; Borek, Hasty, and Tsimring, 2016; Carvalho et al., 2014; Duran-Nebreda and Solé, 2016; Boehm, Grant, and Haseloff, 2018; Cachat et al., 2016). Artificial TPs are expected to have eventual applications in nanotechnology, tissue engineering and regenerative medicine (Scholes and Isalan, 2017; Tan et al., 2018). Despite much effort in this area, engineered TPs remain elusive. Our comprehensive Turing network atlas contains the “blueprints” required for identifying natural TPs, and guiding the engineering of synthetic systems.

## Results

### Developing a tractable model to search for TPs

To generate a network atlas for exploring the design space of TP formation, we need to develop a modeling framework that copes with this computationally demanding task. Our goal is to study the dynamical behavior of spatially-distributed molecule concentrations, and their capability to form stable spatial patterns, across the complete range of 2- and 3-node network topologies. Furthermore, we aim to employ different regulatory functions (competitive and non-competitive Hill functions), over as wide a range of parameters as possible (e.g. diffusion, activation, and repression, etc.) and necessary; we thus forgo mathematical convenience and tackle biologically more realistic models using state-of-the-art computationally intensive, but robust methods.

A network’s capability to form patterns is determined by four factors which can be ordered hierarchically as shown in Figure 1D. First, a network’s topology needs to be defined, that is, the nodes and edges of the network. Next, the functional form of the interactions *f_i_* need to be specified. Subsequently, the parameters of these functions need to be specified. Finally, for a spatial model, we need to describe the molecular diffusion processes.

The first task is to define the network structure (Figure 2A). For this, we only consider connected networks which cannot be split into non-interacting sub-networks, since these would comprise redundant independent sub-networks with smaller node numbers. We further exclude networks with nodes that have no incoming edge, since such nodes do not experience any feedback from the other nodes and will hence always converge to a spatially homogeneous steady state. The influence of such nodes on the rest of the network would thus not have any impact on spatial patterning. Moreover, we exclude networks that have nodes with no outgoing edge as these would purely act as “read-out” modules; they do not feed back to the dynamics of the rest of the network. Finally, we reduce the number of networks using symmetry arguments (Ma et al., 2009; Babtie, Kirk, and Stumpf, 2014), where simply relabeling nodes maps one onto the other. Only one network from such an equivalent group needs to be considered.

**Figure 2:**
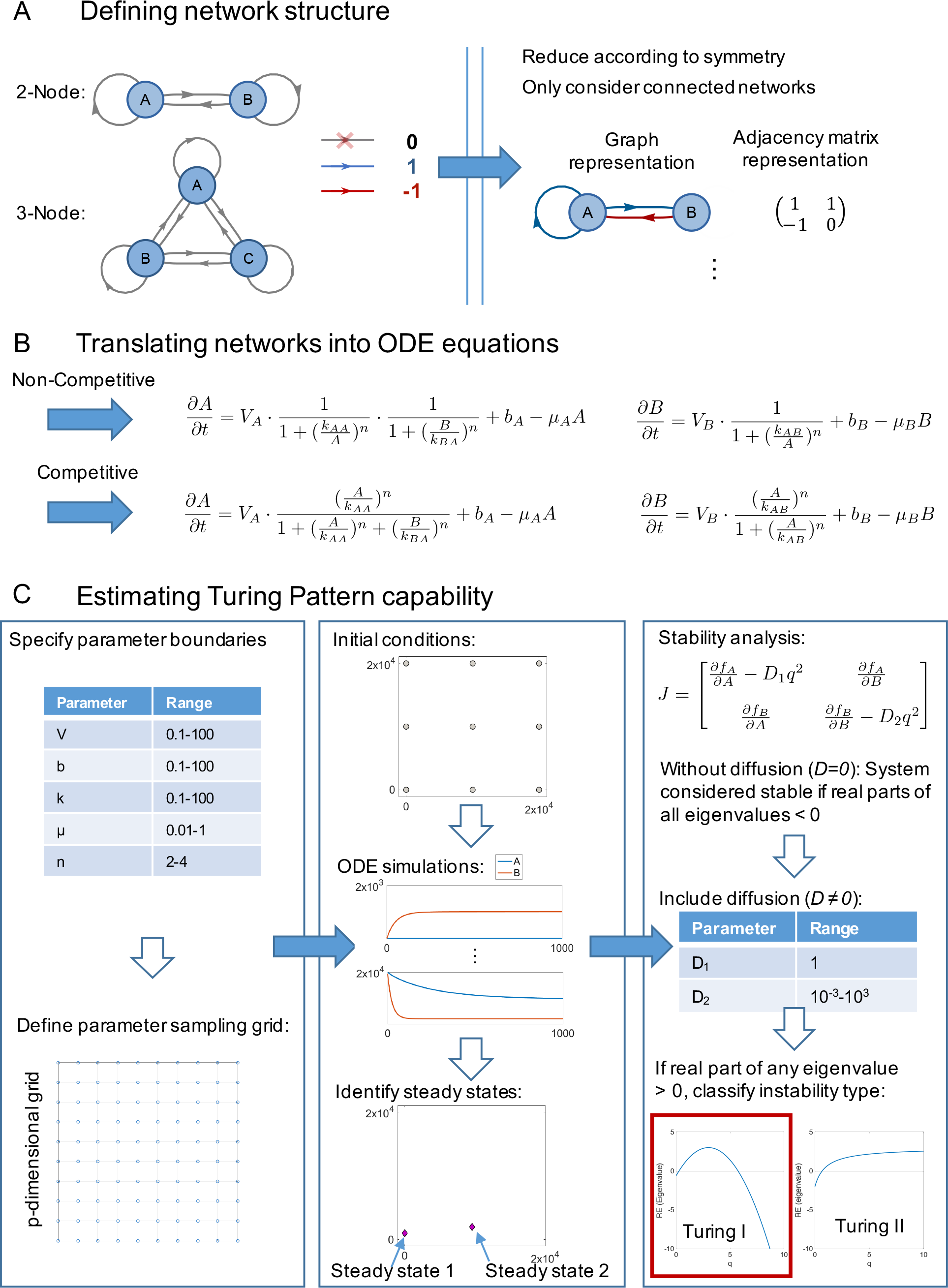
Definition and analysis of networks. (A) Definition of network structure. Left: 2- and 3-node networks containing all possible edges. The latter can be specified as either 0’s (no regulation), 1’s (activation) and −1’s (inhibition). From the set of all possible networks, we identify groups of networks that are equivalent to each other (redundant) and remove all but one network per group. Furthermore, we exclude networks containing any unconnected node(s). The resulting networks can be represented as a network graphs or by their adjacency matrices. The network shown here corresponds to network #8 (see Figure 3A). (B) Generation of ODEs equations. Each edge in a network corresponds to a Hill term in the ODE equations. The way these terms get combined depends on the regulatory mechanism. In the equations, *V* denotes the maximum level of expression and *b* the basal production rate. *n* is the Hill coefficient, indicating the “steepness” of the regulation. We include a linear degradation term with degradation constant *µ* for each species. (C) Workflow for estimating steady states and identifying Turing instabilities. Left panel: for each parameter, a range is specified and a logarithmic grid generated with three values per parameter, amounting to 531441 parameter combinations for fully-connected 3-node networks. Middle panel: for each parameter set, the corresponding ODEs are solved numerically until time *t* = 1000 for several different initial conditions. We use *k*-means clustering on the endpoints of the trajectories to find the steady states of the system. Right panel: the final step of the algorithm calculates the eigenvalues of the Jacobian for each steady state. For the Jacobian evaluated at zero diffusion, the real parts of all eigenvalues are required to be smaller than zero, corresponding to a stable steady state. Subsequently, diffusion is taken into account. A Turing instability exists if the real part of the largest eigenvalues becomes positive for some finite wavenumber *q*. Depending on the behavior of the eigenvalue as a function of the wavenumber *q*, we classify the instability into two types. In this article we only consider Type I instabilities as only these generate patterns with finite wavelengths.

This network pruning amounts a total of 21 and 1934 networks with two and three nodes, respectively. For 3-node systems, we further have to distinguish between which nodes diffuse and which ones are assumed to be stationary. For a system with two diffusing entities this results in 5802 networks (see Supplementary Document S1 for a complete list of network graphs), where A and B denote diffusing nodes and C is assumed to be stationary. Additionally, 1934 networks exist for the case where all three nodes are allowed to diffuse.

Having defined the network topologies, we need to specify the functions and regulatory mechanisms (Figure 2B). In this work, we use Hill-type functions for the regulation as these have been found to fit well with experimental measurements of gene regulatory networks (Estrada et al., 2016; Becskei, Séraphin, and Serrano, 2001; Rosenfeld, Elowitz, and Alon, 2002; Burrill and Silver, 2010; Gardner, Cantor, and Collins, 2000; Ferrell and Machleder, 1998; Ferrell, Tsai, and Yang, 2011; Klumpp, Zhang, and Hwa, 2009). We employ two different types of regulatory mechanisms: fully non-competitive and fully competitive interactions (Figure 2B). For gene regulatory networks, the former corresponds to the situation where all transcription factors independently regulate the corresponding target gene (Figure 1E, left panel). By contrast, for the fully competitive case, all transcription factors compete for their shared or overlapping binding sites (Figure 1E, right panel). In addition to the regulatory interactions, we include a basal production rate and a linear degradation term for each species (see Methods for a detailed description and Figure 2B for an example).

Our approach (Figure 2C) allows us to investigate many different networks and conditions at the appropriate resolution to determine whether they can generate TPs or not. We sample parameters and initial conditions sufficiently densely and are thus able to determine with a high degree of certainty whether a system can exhibit the hallmarks of a TP mechanism: (i) stability of the non-spatial dynamics; with (ii) simultaneous instability of the corresponding spatial dynamics.

Accordingly, to find Turing instabilities, we first need to identify the stable steady states of a given system, and subsequently study their dispersion relation which is defined as the largest eigenvalue of the linearized system as a function of wavenumber *q*. (see Methods for details).

If spatial diffusion is added, it is possible that deviations from the steady state of certain length scales (or, alternatively, wavelengths) do not decay towards the homogeneous steady state, but instead become amplified. This is called a diffusion-driven or Turing instability. If only an intermediate range of length scales experiences such an amplification, we speak of a Turing I instability. In this case, a system typically forms a pattern of the wavelength for which the amplification is maximal (see Figure 1C). Thus, we are able to screen networks and analyze the types of TPs they form. Figure 2C summarizes the computational procedure.

### Turing topologies are common but sensitive to regulatory mechanisms

Analyzing all 2-node topologies (21 networks) and 3-node topologies with two diffusors (5802 networks) we find that more than 60% can exhibit Turing I In-stabilities and thus we expect them to be capable of generating TPs (Figure 3). This large number of potential Turing networks is many fold higher than the number of networks identified in the literature to date (≈ 700 topologies) (Zheng, Shao, and Ouyang, 2016; Marcon et al., 2016).

**Figure 3:**
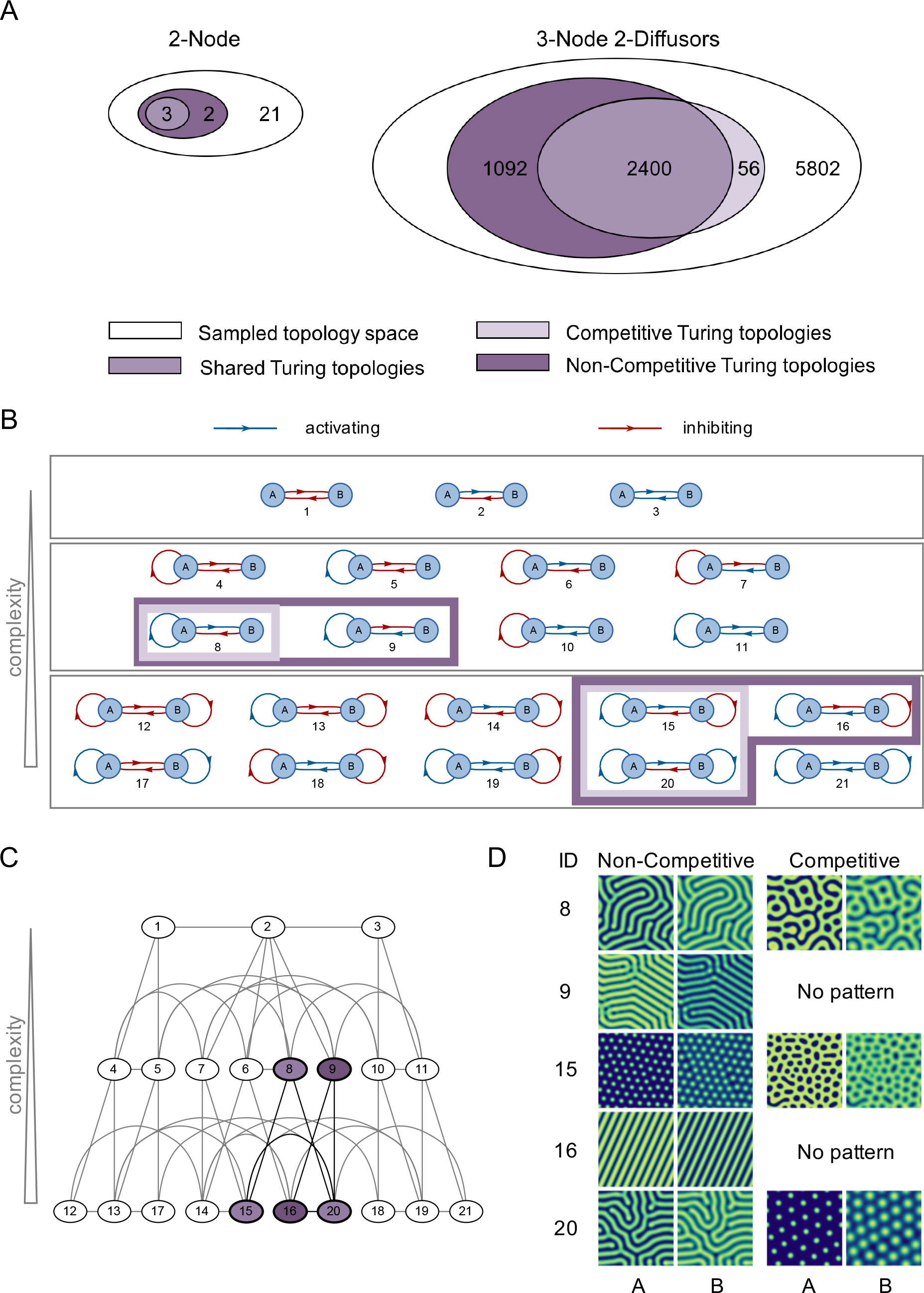
Results for 2-node networks. (A) Visualization of the sampled network topology space and fractions of Turing pattern generators (Turing topologies) for 2- and 3-node networks with 2 diffusing molecules. (B) Considered 2-node networks selected according to criteria described in Figure 2A. Blue and red edges indicate activation and inhibition, respectively. The networks are arranged according to their complexity (that is, the number of edges) with increasing complexity towards the bottom. Each network is given an ID number (1-21). Lilac boxes indicate the networks identified as Turing pattern generators. Dark lilac indicates non-competitive regulatory mechanisms, whereas the lighter shade indicates competitive ones. (C) Hierarchical graph of 2-node networks. Each node represents a network of a given ID number. Networks are connected by an edge whenever they can be transformed into each other by addition, deletion or modification (change of sign) of a single edge. For example network #1 can be transformed into network #4 by addition of a negative self-interaction on node A, and network #1 into network #2 by changing the sign of one edge from inhibition to activation (see networks in (B)). The nodes are colored according to the legend shown in (A), with different shades of lilac indicating for which regulatory mechanism the networks exhibit Turing I instabilities. (D) Exemplary 2-dimensional patterns for the identified 2-node Turing generating networks. The parameters for which the patterns were generated are given in Supplementary Document S3.

Importantly, we observe that subtle features beyond network structure influence a network’s pattern generating capability. The first difference appears in the choice of regulatory mechanism. We find that there are fewer competitive Turing topologies overall among the 2-node networks (Figure 3A). All five networks are detected for non-competitive mechanisms, of which only three are found for competitive interactions (Figure 3B). However, we find that the rarer competitive interactions are more robust to parameter variations than non-competitive interactions, which results in more TP solutions within a given topology. Network #8 constitutes the classical Turing network, which was analyzed by Alan Turing in 1952 (Turing, 1952) (Figure 3B-D). The other four 2-node networks have also been reported elsewhere (Zheng, Shao, and Ouyang, 2016; Marcon et al., 2016).

For 3-node systems, we similarly find a group of Turing topologies that is shared by both regulatory mechanisms (2400 networks, Figure 3A). While a large fraction of topologies exhibit a Turing I instability, this is again mainly for non-competitive rather than competitive interactions. This result stands in sharp contrast to the existing literature which mainly highlights the network topology as the important factor for TP capability (Diego et al., 2017). By contrast, our results suggest that network structure alone does not suffice but that the choice of regulatory function also critically determines a network’s Turing capability.

### Minimal topologies define key properties such as pattern phasing

In order to determine key dynamic features of networks it has been shown to be advantageous to analyze the minimal topologies necessary to achieve a particular behavior. For example, such an analysis revealed that the key motifs to achieve single stripe patterns mediated by external cues (French Flag patterns) are incoherent feed-forward loops (Ingolia and Murray, 2004; Schaerli et al., 2014; Cotterell and Sharpe, 2010).

We perform a similar analysis here to identify minimal TP motifs. To this end, we create a network atlas in which all networks that differ by a single edge are connected (see atlas for 2-node systems in Figure 3B,C). Each connection between a network represents the addition/deletion of a single edge, or a change of sign of a single edge. Networks are subsequently sorted hierarchically according to their complexity (here defined as network size). From all networks that can generate TPs, we identify two minimal or “core topologies” for 2-node networks: #8 for the competitive case and #8 and #9 for the non-competitive case. All other Turing networks can be constructed from these by the addition of one or more edges.

Turing networks can show patterns in which the concentration maxima of the different molecular species are either in phase or out of phase. “In phase” refers to systems in which the maximal concentrations of both species coincide, whereas for out of phase mechanisms, the maximum of one species coincides with the minimum of the other (see Figure 4D). We analyzed the 2-node Turing topologies (given in Figure 3B) with respect to their patterns’ phases, by numerically solving the corresponding PDEs. We find that all 2-node Turing networks with competitive regulations give rise to in-phase patterns. In the competitive case, it appears that the networks #15 and #20 inherit the patterning phase from the core network #8, to which they can be reduced.

**Figure 4:**
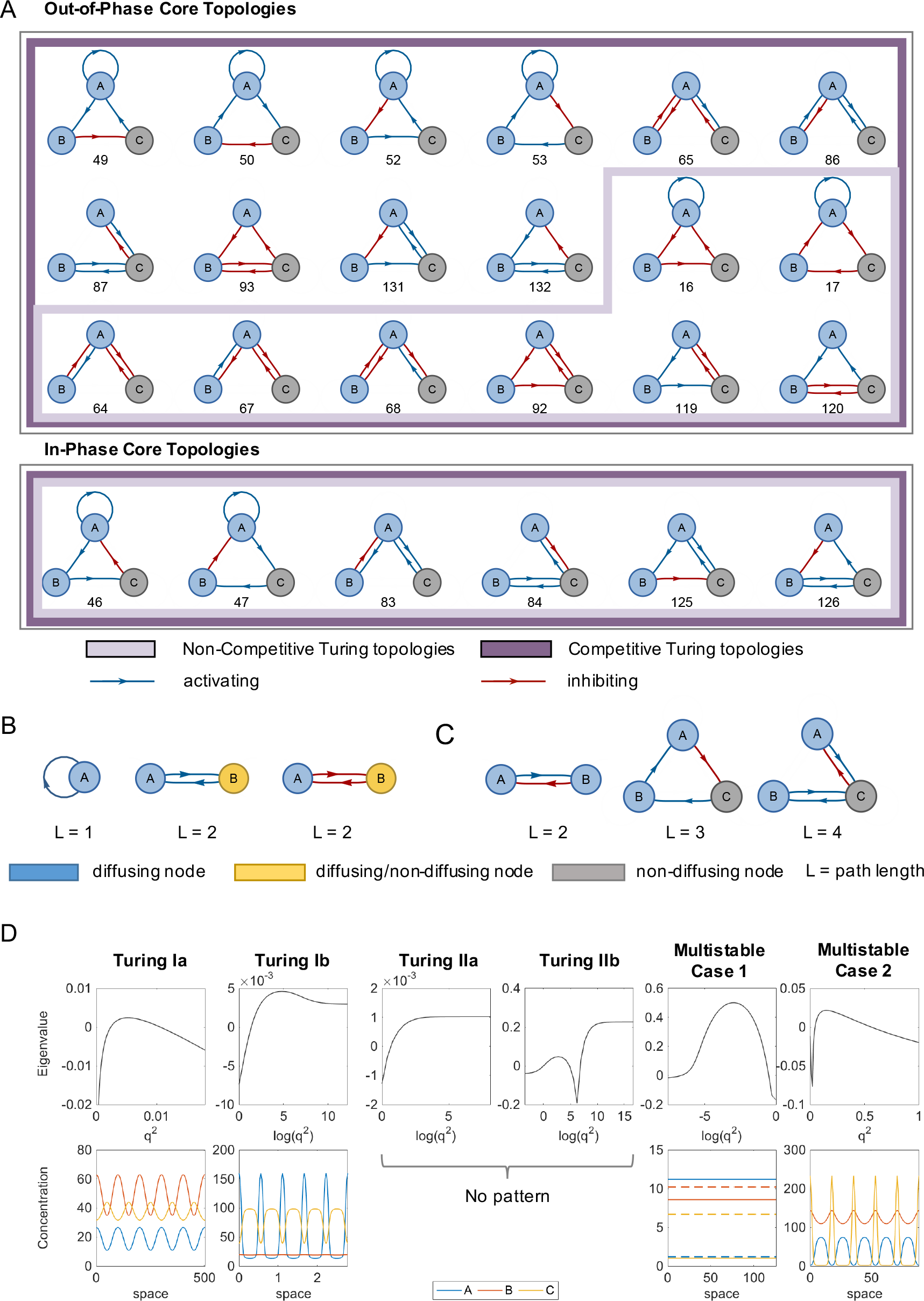
Core networks and core motifs for 3-node networks. (A) Top: 3-node core topologies that generate out-of-phase patterns (one species’ maximum concentration is shifted by half a period with respect to the other two). Bottom: 3-node core networks exhibiting in-phase patterns (all concentration maxima aligned). Colored frames indicate the regulatory mechanism (competitiveness) for which the networks exhibit Turing I instabilities. (B) Examples for the first identified core motif: positive feedback loop of length one or two on one of the diffusing nodes. The figure shows the different possibilities to achieve this. If the path length is two it can be mediated by either the other diffusing or the stationary node. (C) Examples for the second identified core motif: diffusion-mediated negative feedback on a diffusing node. One diffusing node has to have a negative feedback loop whose path includes the other diffusing node. This interaction can consist of two, three or four edges and the figure shows one example each. The core motifs in (B) and (C) are almost sufficient and almost necessary for TPs. (D) Examples of different Turing instabilities and resulting patterns. Note that for Turing IIa and Turing IIb instabilities no patterns are formed. Note also that in the multistable case 1, no pattern is generated despite the existence of a Turing I instability. Instead, starting from a perturbation around the stable steady state with Turing I instability (indicated as dashed lines) this moves to a second homogeneous steady state. This behavior is observed for 4% (non-competitive) and 14 % (competitive) of multistable network-parameter combinations exhibiting a Turing I instability. The parameters for which the dispersion relations and patterns were generated are given in Supplementary Document S3.

For non-competitive regulation, network #8 again exhibits only in-phase patterning, whereas network #9, which constitutes the second core network for non-competitive systems, shows out-of-phase patterning. One might thus expect network #20 (which can be reduced to either network #8 or network #9 by removal of one edge) to give rise to both in and out-of-phase patterns. Our analysis shows that this is in-deed the case. Interestingly, here the phase can be controlled by the diffusion constants: when A diffuses faster than B, in-phase patterning is observed, whereas if B diffuses faster than A, out-of-phase patterning is seen. In a sense, the choice of which node contains the “fast” or “slow” species thus becomes a topology-related system parameter. Overall, key qualitative properties of TPs such as phasing appear to be mediated by the underlying core topologies.

### Core topologies also specify phase properties for 3-node networks

We next apply the complexity atlas reduction procedure to the 5802 different 3-node networks where two species can diffuse (3N2D). As for the 2-node networks, we reduce 3-node networks to their core topologies. This leads to a hierarchical graph similar to the 2-node case shown in Figure 3B, but that can-not be depicted due to its large size (> 2400 nodes and > 10^4^ dependencies). Within this atlas, we find 12 and 20 core topologies for competitive and non-competitive regulation, respectively, all of which have four edges (Figure 4A). As in the 2-node case, the competitive core topologies constitute a subset of the non-competitive systems. We also assess whether these core topologies give rise to in phase or out-of-phase patterns, by solving the corresponding PDEs numerically. In phase mechanisms (Figure 4A bottom) are observed less commonly than out of phase mechanisms (Figure 4A top). One might expect that in phase mechanisms are rarer with three nodes, since more species now have to be in phase. In contrast to the 2-node case, however, some competitive systems now also form out-of-phase patterns. In all core topologies, we solely observed either out of phase or in-phase patterns, which suggests that these minimal topologies exhibit unique behaviors.

Analyzing 3-node networks with three diffusing nodes, we find that all such 3N3D Turing networks are also a 3N2D Turing network. We further find that the core topologies for two and three diffusing molecules coincide. This suggests that networks in which three molecules diffuse are mere expansions of the Turing parameter set, but that the instability driving component already exists in the two-diffuser systems. Therefore, in order to understand the core TP mechanisms for 3-node systems, we have to focus on those with two diffusing species.

### Two core motifs account for more than 95% of Turing topology space

Analyzing all 3-node core topologies with Turing I instabilities, we identify two frequently occurring core motifs: a positive feedback on at least one of the diffusing nodes, consisting of one or two edges (Figure 4B), and a diffusion-mediated negative feedback loop on both diffusing nodes (Figure 4C).

For the first core motif, the positive feedback loop, three possible configurations exist: a direct positive feedback (e.g. network #49) or an indirect positive feedback consisting of either two positive or two negative edges to another node. The interaction can either be mediated via the other diffusing node (e.g. network #65) or the non-diffusing node (e.g. network #131). The second core motif consists of a negative feedback loop on one of the diffusing nodes that is mediated through the other diffusing node. This motif can be of several different types and some examples are depicted in Figure 4C.

Having identified the two core motifs from studying the 3-node core topologies, we re-analyzed all 2-node networks with respect to these motifs. We find that all 2-node Turing networks do indeed possess both core motifs, and all networks containing both motifs exhibit Type I instabilities in the non-competitive case (Figure 3A). For the 3-node networks, we find that 96% of all networks containing both core motifs do exhibit Turing I instabilities. Since we can only sample a finite number of parameters, we cannot categorically rule out that the missing 4% might also exhibit a Turing I instability for certain parameters. To reduce the probability of missing Turing I instabilities for these networks, we ran additional simulations for 10^5^ randomly sampled parameter sets per network. Analying all networks exhibiting Turing I instabilities, we observe that 95% possess both core motifs. We confirm by simulation that the remaining 5% topologies do indeed give rise to TPs despite the absence of the core motifs. Therefore, the two identified core motifs together constitute an almost necessary and almost sufficient criterion for Turing I instabilities.

### Differentiating Turing Instabilities - new instability types and their patterns

The dispersion relation of a system is typically related to the resulting pattern: the wavenumber *q*_max_ for which the dispersion relation assumes a global maximum (that is, the largest eigenvalue of the Jacobian becomes maximal; see Methods) experiences the largest amplification. We thus expect to see a pattern with wavelength 2*π*/*q*_max_ (see Figure 1C).

Due to the extent of this analysis in terms of the number of networks considered, we also observe some qualitatively novel types of dispersion relations that had not been reported previously in the literature. We distinguish four different groups of dispersion relations. First, the “classical” Turing I instability fulfills three criteria: for *q* = 0, the system should be stable (i.e. Re(λ) < 0); for a finite *q* we have Re(λ) > 0; and Re(λ) < 0 for *q* ⟶ ∞. This type of instability is the most commonly discussed mechanism underlying TPs. We describe as a Turing II instability the case when the steady state is stable for *q* = 0 (i.e. Re(λ) < 0), where there exists some threshold *q*\* such that Re(λ) > 0 for all *q* > *q*\*, and the dispersion relation becomes maximal for *q* ⟶ ∞. Therefore, for *q* > *q*\*, the larger the mode *q* the stronger the amplification. Consequently, arbitrarily large modes and hence short wavelengths get amplifyed the most, which does not lead to a stable pattern with a well-defined wavelength.

The previous cases are referred to as Type Ia and IIa instabilities. However, other types of dispersion relations exist for diffusion driven instabilities that do not fall into either Type Ia or IIa. Figure 4D shows examples for all four categories that we find. One of these is similar to Type Ia and capable of producing stable patterns and we hence term it Type Ib. The other dispersion relations does not produce patterns and we denote it by IIb. Type Ib instabilities fulfill both criteria of a negative real λ for *q* = 0 and possess a finite *q*_max_ for which the dispersion relation becomes positive and assumes a global maximum. However, the dispersion relation does not fulfill the third criteria of Re(λ) < 0 for *q* ⟶ ∞, but remains positive instead. Since the dispersion relation does possess a global maximum in *q*_max_ and hence a finite wavelength that experiences the strongest amplification, one might expect such systems to lead to patterns anyway. Through numerical simulations we verify that this is indeed the case. This result agrees with the conclusion of a recent preprint (Smith and Dalchau, 2018). An exemplary pattern plot is visualized below the corresponding dispersion relation in Figure 4D.

Figure 4D panel 4 shows the Type IIb dispersion relation. Similarly to the Type IIa instability, the dispersion relation becomes positive for some finite *q*_1_ and assumes its maximum for *q* ⟶ ∞. However, in contrast to the original Type IIa case, the dispersion relation becomes negative again for some intermediate *q*_2_ > *q*_1_; but like the Type IIa case the maximum of this dispersion relation occurs for *q* ⟶ ∞ and thus we cannot get stable patterns since arbitrarily small wavelengths experience the strongest amplification. To our knowledge this type of Turing instability has not been reported in the literature before.

We would like to note that in the analyses described in the previous sections we merely distinguished systems according to their patterning capability and referred to Ia and Ib jointly as “Type I”, and IIa and IIb jointly as “Type II”. Both types of instabilities are important when considering the analysis of reallife systems. Ib could be mistaken for a non-patterning system as it does not fulfill a classical criterion for a Turing I instability, which may lead to underestimating the robustness of a system. Similarly, IIb can be mistaken for a Type I instability (if *q* is sampled over an insufficient scale) consequently leading to an overestimation of Turing robustness.

### Turing I instabilies are not a sufficient criteria for patterning

In development, cell-fate decisions are often governed by systems in which multiple stable steady states exist (Harrington et al., 2014; Harrington et al., 2013; Moris, Pina, and Martinez Arias, 2016). In mammalian stem cells, for example, Nanog, Sox2 and Oct4 form a system possessing two stable steady states. Here, each steady state corresponds to one specific cell fate: either a cell remains pluripotent or it differentiates.

We therefore must explore how multi-stability affects TP formation. First, we find that systems with multiple stable steady states can exhibit Type I instabilities (Ia or Ib), either for a single or for several steady states. The former has also been reported in a recent preprint (Smith and Dalchau, 2018). Similarly, when probing the patterning behavior of Type Ia, Ib, IIa and IIb instabilities, we simulate the spatial system numerically for all systems and parameter combinations for which either one or multiple Turing I instabilities are present. Even though most systems indeed show the expected pattern formation (86% and 96 %, for competitive and non-competitive systems, respectively), some do not (14% for competitive and 4% for non-competitive). Rather than observing stable, reproducible patterns, we find that these systems transition from a perturbation around the steady state with a Turing I instability to one of the other stable steady states. Figure 4D, Panel 5, shows an example of such behavior. We thus conclude that a Type I Turing instability is not a sufficient criterion for pattern formation in multi-stable systems. Again this highlights the need for a thorough analysis if we want to be able to predict and validate natural TP generating networks.

### Defining quantifiable measures of robustness

In biological systems, parameter values are generally known only with partial certainty, and sometimes entire interactions are unknown (Kirk, Babtie, and Stumpf, 2015). It is therefore crucial not only to identify networks that exhibit Turing instabilities, but also to assess their sensitivity with respect to these uncertainties.

To this end, we define four measures of robusness to parameter variation or uncertainty: robustness to intracellular processes, robustness to extracellular processes, topological robustness and total robustness. As intracellular parameters, we define all kinetic parameters of the ODE equations (Figure 2B) that describe the chemical interactions between species within cells, and we define “intracellular robustness” as the fraction of the analyzed parameter combinations that is capable of Turing pattern formation.

In addition to intracellular processes, the speed of extracellular diffusion of molecules determines if a network possesses a Turing instability. We accordingly define the "extracellular robustness" as the robustness of a Turing network to changes in the diffusion constants, given that the intracellular parameters are fixed to values that can give rise to a Turing instability (see Figure 5A).

**Figure 5:**
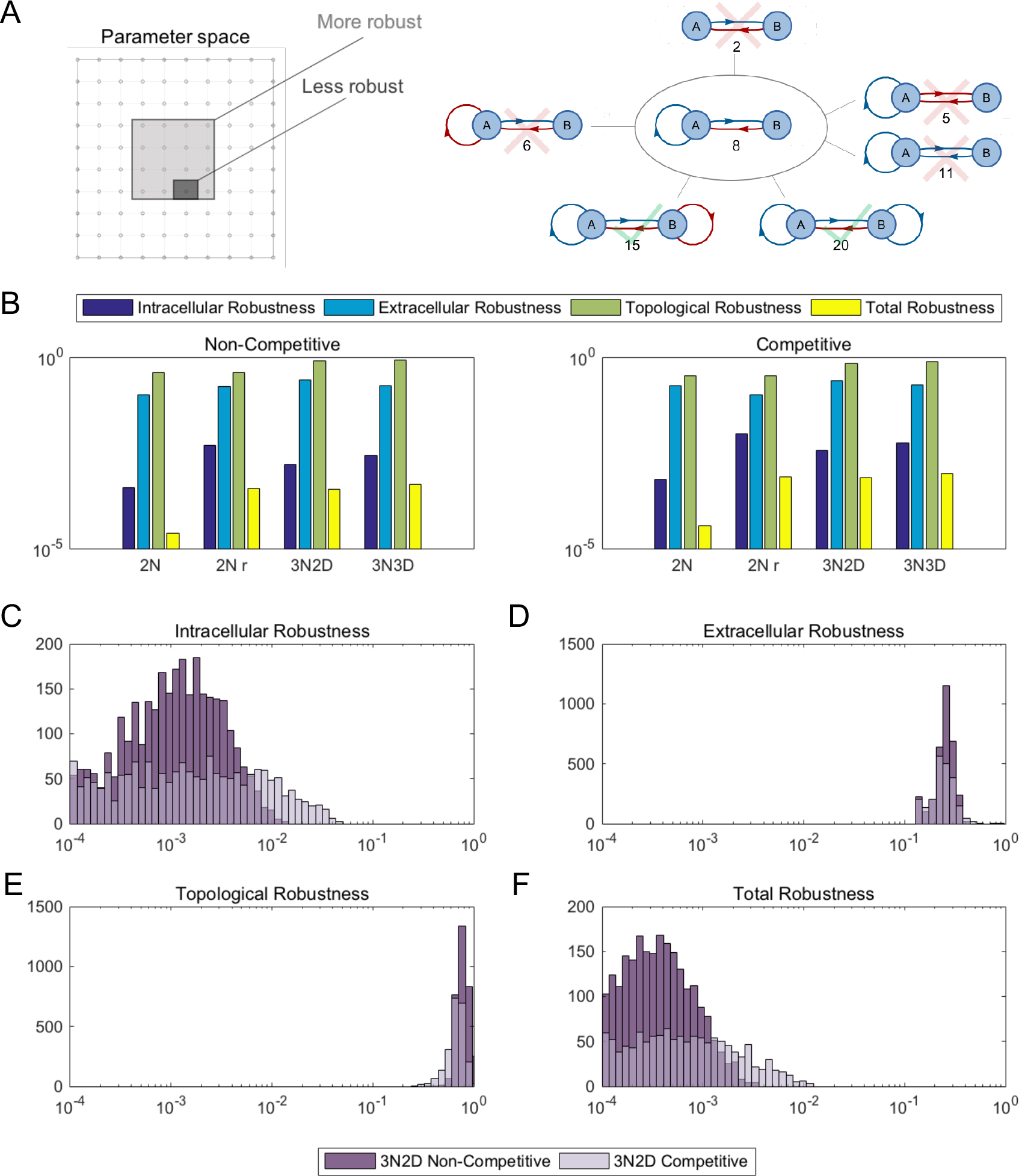
Quantifying a network’s Turing capability: defining four measures of robustness. (A) Left: definition of intracellular (extracellular) robustness as the fraction of sampled kinetic (diffusion) parameter sets leading to Turing I instabilities. Right: definition of topological robustness as the fraction of neighboring networks (that is, networks into which a given network can be transformed by addition, deletion or modification of a single edge) that exhibit Turing I instabilities. The example shows that two out of six neighbors of network #8 are Turing networks, leading to a robustness 1/3. The total robustness is defined as the product of intracellular, extracellular and topological robustness. (B) Mean values for the different robustness measures for competitive (left) and non-competitive systems. 2*N* (2Nr) denotes 2-node systems for which V and b are varied (restricted to 100 and 0.1, respectively). 3N2D and 3N3D represent 3-node systems with two and three diffusing nodes, respectively. (C-F) Histogram plots of the four robustness measures comparing for both regulatory mechanisms. We find that competitive systems are on average more robust for intracellular processes, whereas non-competitive systems are topologically more robust. In terms of total robustness, a subset of 3N2D competitive networks constitute the best performing networks.

Despite uncertainties in parameter values, it is frequently the case in biological experiments that one cannot even be certain about the network topology of a given system (Babtie, Kirk, and Stumpf, 2014). There might be additional, unknown regulatory interactions between species, or assumed interactions might not be active. Accordingly we define “topological robustness” as follows: for a given network, consider all networks that can be generated by adding, removing or changing one edge (as exemplified in Figure 3C for 2-node systems). Then the topological robustness is defined as the fraction of generated networks that are capable of exhibiting Turing I instabilities (Figure 5A).

Finally, we would like a measure for the overall robustness of a given network taking into account the influence of all three different sources of uncertainties. This “total robustness” is the product of the intracellular, extracellular and topological robustness (Figure 5).

### Competitive 3-node systems are the most robust Turing networks

We compute the intracellular, extracellular, topological and total robustness for all 2-node and 3-node networks (Figure 5). We find that Turing networks with competitive interactions are more robust than non-competitive ones, in particular with respect to intracellular parameters. This is consistent for 2-node systems (~ 1.7-fold), as well as 3-node systems with two (~ 2.3-fold) and three (~ 2.1-fold) diffusing species (see Figure 5B). As the intracellular robustness varies over about two orders of magnitude more than the extracellular and topological robustness, competitive systems are also in total more robust (~ 1.5 – 2-fold). Our results demonstrate that the choice of regulatory interactions can have significant influence on a network’s Turing capability: non-competitive topologies are more likely to be able to generate TPs whereas competitive systems that do generate TPs are more robust.

Due to computational cost, the *V* and *b* parameters were restricted for the analysis of 3-node systems (see Methods). Consequently, to compare like-for-like, we calculated the robustness for 2-node systems under the condition that *V* and *b* were fixed to the same values. With this restriction on parameter space, 2-node systems are on average more robust to intracellular variations than 3-node systems (~ 2.7-fold for competitive and ~ 3-fold for non-competitive regulations). On the other hand, 3-node systems are on average significantly more robust to extracellular (~ 2.4 (competitive) and ~ 1.5-fold (non-competitive) and topological (~ 2-fold) variations than 2-node systems. Even though in total the average robustness of 2 compared to 3-nodes is not significantly different, we do find that amongst the top (most robust) networks for either case, 3-node systems are more than 4-fold more robust in total than the top 2-node systems. It is thus likely that TP networks in nature will be composed of at least three interacting species.

### Robustness maps of 3N2D topology space to reveal the most robust networks and their neighborhoods

Due to the large number of 3-node Turing topologies we visualize their total robustness in a “robustness map” shown in Figures 6A (non-competitive) and 7A (competitive). We group networks according to their complexity; for each complexity class we additionally depict the most robust network (right panel). However, these networks only constitute a fairly small fraction of the top 10 most robust networks (three and two for competitive and non-competitive, respectively; Figures 6B and 7B). Overall, networks with complexity of 5-6 (competitive) and 6-7 (noncompetitive) are the most robust topology groups. We find no significant correlation between robustness values and topology complexity. This further suggests that increasing complexity does not generally lead to a larger robustness. All robustness measures (intracellular, extracellular, topological and total robustness) which we derived from this analysis are provided in Supplementary Document S2; network identifiers correspond to those in the network graphs provided in Supplementary Document S1.

**Figure 6:**
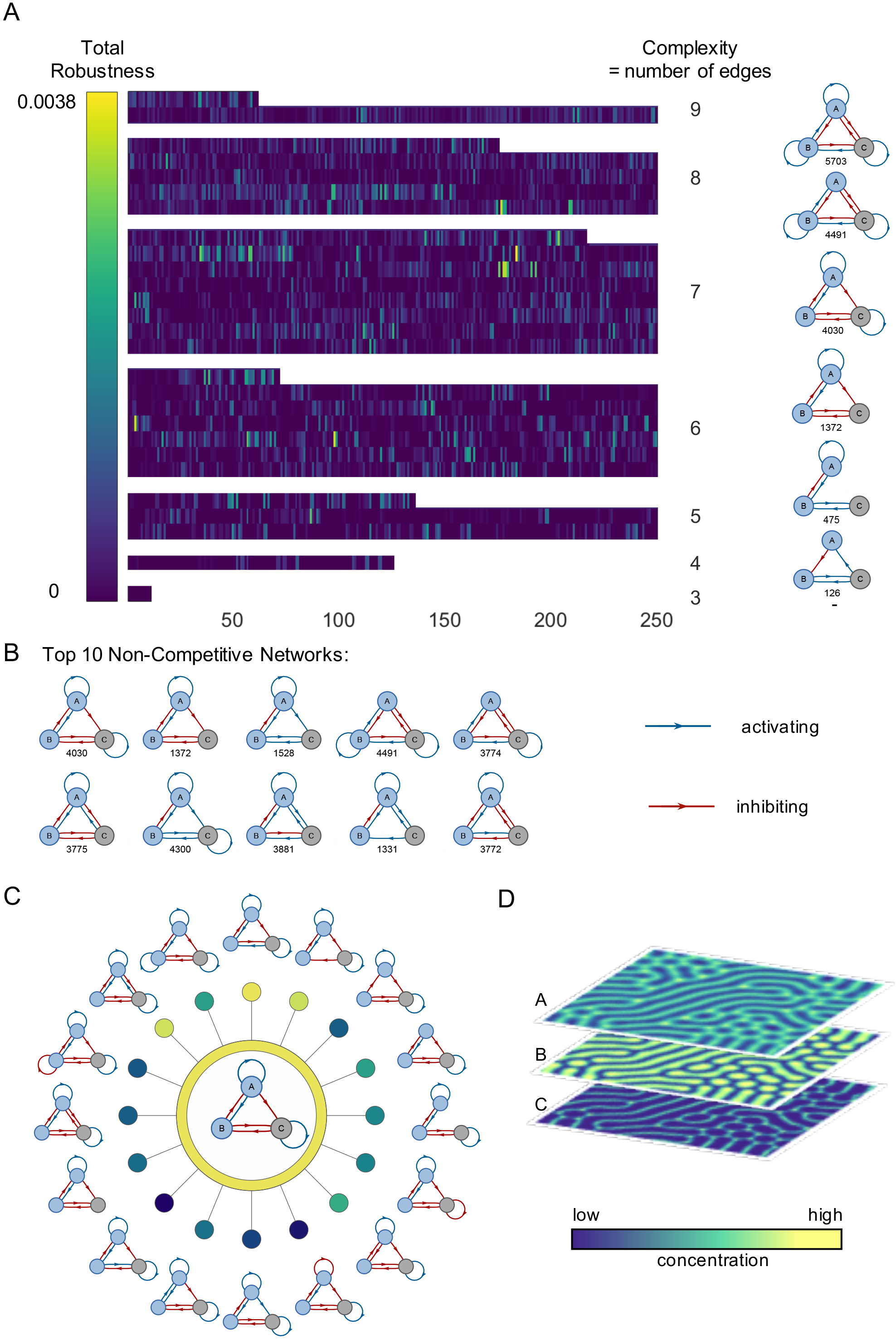
Most robust 3-node Turing networks with non-competitive regulation and two diffusing nodes. (A) Robustness map. The figure indicates the total robustness of all analyzed 3-node networks arranged according to complexity. For each complexity class we show the most robust network, together with its ID number. (B) Ten most robust networks with numbers indicating the corresponding network ID. Total robustness decreases towards the right and bottom, i.e. #4030 is the most robust network. The absolute robustness values are given in Supplementary Document S2. (C) Local neighborhood atlas for the most robust non-competitive Turing network #4030 (center). Colored nodes indicate the total robustness value according to the colorscale shown in (A). (D) 2-dimensional pattern for the most robust non-competitive Turing topology #4030. Yellow and blue indicate high and low species concentrations, respectively, for nodes A, B and C. This is an out-of-phase pattern. The parameters for which the pattern was generated are given in Supplementary Document S3.

Due to the large number of topologies, we are not able to illustrate the full network atlas to show how these networks are related. Instead, we provide a local neighborhood atlas for the single most robust noncompetitive and competitive networks (Figure 6C and 7C) and a corresponding 2D PDE solution (Figure 6D and 7D). The local neighborhood atlas contains all the networks that can be generated from the central network by adding, deleting or modifying one edge. Most edge changes lead to a pronounced drop in Turing robustness, although it is still possible for evolution to “walk” from one topology to another while still maintaining a TP.

Overall, one of the most important conclusions from this study is that Turing network topologies and mechanisms are more common within random networks than previously thought. Therefore, it is possible that evolution can move through these less robust — but relatively common — prototypic solutions, towards one of the more robust networks.

**Figure 7:**
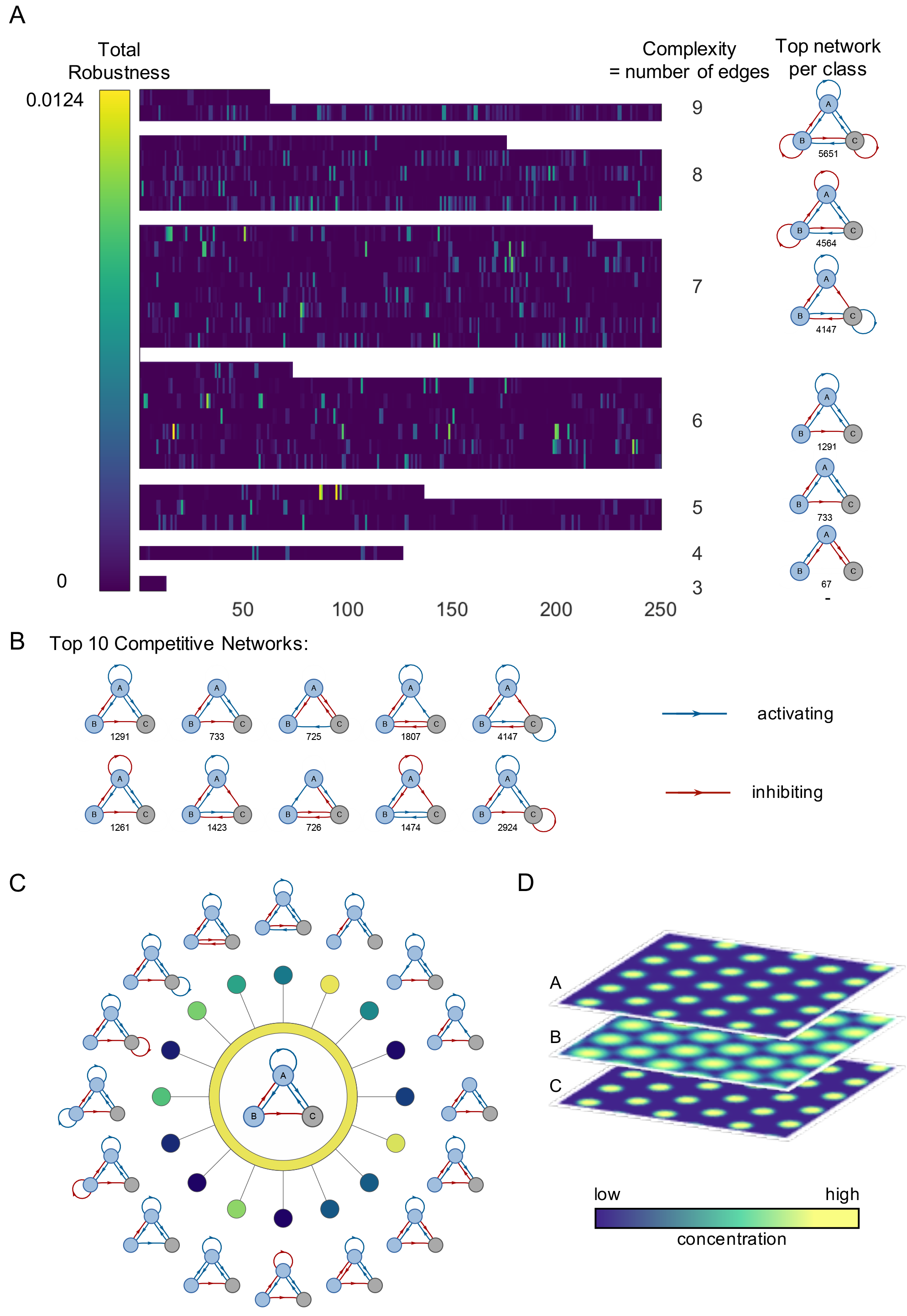
Most robust 3-node Turing networks with competitive regulation and two diffusing nodes. (A) Robustness map. The figure indicates the total robustness of all analyzed 3-node networks arranged according to complexity. For each complexity class we show the most robust network, together with its ID number. (B) Ten most robust networks with numbers indicating the corresponding network ID. Total robustness decreases towards the right and bottom, i.e. #1291 is the most robust network. The absolute robustness values are given in Supplementary Document S2. (C) neighborhood atlas for the most robust competitive Turing network #1291 (center). Colored nodes indicate the total robustness value according to the colorscale shown in (A). (D) 2-dimensional pattern for the the most robust competitive Turing topology #1291. Yellow and blue indicate high and low species concentrations, respectively, for nodes A, B and C. This is an in-phase pattern. The parameters for which the pattern was generated are given in Supplementary Document S3.

## Discussion

Turing’s pattern generating mechanism, later independently rediscovered by Gierer and Meinhardt, is an elegant way in which purely biochemical mechanisms can give rise to reproducible and self-organizing spatial patterns. Despite initial unease (and sometimes outright hostility) over them being relevant and robust mechanisms of patterning, TPs are now widely accepted and have become an important cornerstone of modern developmental biology.

Given their provenance, it is perhaps not surprising that TPs have also received close attention by mathematical modelers interested in biological pattern formation. But these have typically focused on single models, exploring them in great detail. The largescale *in silico* surveying of potential TP models is a much more recent phenomenon. There are two potential pitfalls in such analyses: (i) computational cost may require simplified models or prohibit exhaustive analysis; (ii) automating any mathematical analysis, but in particular something as subtle as pattern formation is non-trivial unless we have very precise criteria by which stability, robustness and patterns can be scored. Our approach addressed these issues explicitly and from the outset, and the algorithm employed here is capable of analyzing a wide range of different network structures and is independent of functional choices for regulatory mechanisms and rate functions.

The rewards of a thorough and comprehensive analysis of 2-node and 3-node candidate Turing network models are also considerable. This analysis clearly demonstrates that the structure of the network alone cannot determine whether a TP exists; this is in line with a large body of work on network dynamics (Ingram, Stumpf, and Stark, 2006), but seems to directly contradict results from other studies on TP mechanisms. These, however, (i) fail to fully assess the dependence of TP on regulatory mechanisms and reaction rate parameters (largely because a mathematically more convenient model structure was imposed); and (ii) their models are special cases of the more general and more comprehensive treatment here.

Perhaps the most surprising result of this analysis is how common networks capable of producing TPs are. Evolution has had many opportunities to ‘stumble’ across a Turing network (more than 60% of networks considered here can produce TPs), even though for most architectures the existence of a TP depends crucially on parameters. Being common but not very robust to parametric and structural changes could suggest that many different architectures are used in nature to generate TPs (as was already hinted at in some of Meinhardt’s work (Meinhardt, 2013)). Once a structure is in place with suitable regulatory interactions and reaction rate parameters, and the resulting TP confers an evolutionary advantage, natural selection is likely to maintain this mechanism.

While network structure by itself neither guarantees nor implies the existence of a TP, the overwhelming majority of TP generating mechanisms embed the hallmarks encapsulated by two core motifs: a positive feedback on at least one of the diffusing nodes and a diffusion-mediated negative feedback loop on both diffusing nodes. It is tempting to speculate that larger systems will also reflect these compositional rules and have these core motifs embedded; a basic TP generating motif could thus, for example, be regulated in a more nuanced manner.

Finally, our exhaustive analysis reveals a spectrum of Turing-like instabilities, and subtle dependencies between these and the eventual pattern formation. These are naturally easily missed when analyzing individual Turing systems, or applying simplifying measures in large-scale surveys.

The analysis presented here provides us with a set of blueprints for TP generating mechanisms that can guide the search for naturally evolved Turing systems, as well as the *de novo* engineering of biosynthetic systems. For the latter, in particular, competitive networks should be a safer bet, due to their increased (intracellular) parametric robustness. This tension between commonality and robustness is perhaps one of the most fascinating (or vexing) features of Turing mechanisms.

## Methods

### A Semi-Formal Summary of our TP search

This first section provides an overview of the computational approach taken here; more technical detail is provided in the following sections; with this information in hand it should be straightforward to implement and repeat our analysis (software is made available, below, too).

To identify potential Turing instabilities, we first consider the stability of the non-spatial system. Suppose we have a system of *n* interacting molecular species with time-dependent concentrations *x*_*i*_(*t*), *i* = 1, …, *n*. We model the dynamical behavior of such concentrations in terms of a system of ordinary differential equations (ODEs),

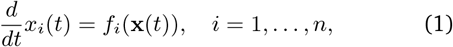

where **x**(*t*) = (*x*_1_(*t*), …, *x*_*n*_(*t*)), *x*_*i*_(*t*) ∈ ℝ is the concentration of the *i*th species at time *t*. Equations of the form in cannot typically be solved exactly for nonlinear functions *f*_*i*_. However, efficient numerical algorithms exist to solve such equations approximately, leading to time trajectories of the system, that is, to solutions *x*_*i*_(*t*) (see Figure 2C for examples). **f**(**x**(*t*)) = (*f*_1_(**x**(*t*)), …, *f*_*N*_(**x**(*t*))) in equation (1) encodes the interactions between the different species. If the *xi* denote protein concentrations, for example, **f**(**x**(*t*)) may encode the regulatory mechanisms between the proteins.

Figure 1A shows a network representation of the Gierer-Meinhardt model, and its governing ODEs (Gierer and Meinhardt, 1972). The nodes indicate the interacting species and arrows the direction of interaction. We distinguish between two possible types of interactions: activating (blue arrows) and inhibiting (red arrows), which we encode in the corresponding components of **f**(**x**(*t*)), whose functional form is not specified by the graph (we employ Hill functions in the reaction equations).

We next include the spatial diffusion of molecules and extend the model in Equation (1) to a spatial setting. In this case the molecule concentrations *x*_*i*_(*t*) become space-dependent concentration fields *x*_*i*_(*r*, *t*), where *r* denotes the spatial location. These fields satisfy the set of coupled partial differential equations (PDEs)

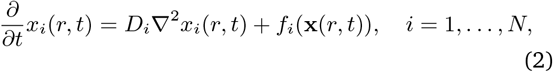

which is obtained from Equation (1) by adding the diffusion term *D*_*i*_ **∇**^2^*x*_*i*_(*r*, *t*), where *D*_*i*_ is the diffusion constant of the *i*th species and **∇** the gradient with respect to position *r*, and concentration is now a function of space and time, *x*_*i*_(*r*, *t*).

The next step is to screen for the formation of stable patterns through diffusion-driven instabilities (Maini et al., 2012). In this case, for any small spatial fluctuations one may expect the concentrations *x*_*i*_(*r*, *t*) to become spatially constant again for large times. For many systems, this is indeed the case, but for some systems the interplay of diffusion and reactions can lead to the molecules’ concentrations forming spatial patterns with certain wavelengths that are stable and reproducible in time.

In the Turing pattern framework we start with, **x**\*, which is the stable steady state concentration of the non-spatial system in (1), i.e. *f*_*i*_(**x**\*) = 0, *i* = 1, …, *n* in Equation (1): if the system is in state **x**\*, it remains there for all times, and if the system is close to **x**\* it will converge towards **x**\*. If spatial diffusion of molecules is included into the model, as described Equation (2), it is possible that deviations from the steady state of certain length scales do not decay towards the homogeneous steady state, but are instead amplified. This is then the diffusion-driven or Turing instability. If only an intermediate range of length scales experiences such an amplification, we speak of a Turing I instability. In this case, a system typically forms a pattern of the wavelength for which the amplification is maximal (see Figure 1C).

Mathematically, the stability is determined by the dependence of the real part of the largest eigenvalue of the Jacobian matrix of the system on the wavenumber *q* (see below for more details). We also call this dependence the dispersion relation. For zero wavenumber, a spatially constant system, the real part of the largest eigenvalue is negative, if evaluated at a stable steady state of the non-spatial system. If, however, the dispersion relation becomes positive for some wavenumber, these become amplified and the steady state unstable. For cases where the dispersion relation has a maximum for a finite value of *q*, a Turing I instability is present. If the dispersion relation remains positive and becomes maximal for large wavenumbers, we speak of a Turing II instability. In this case deviations on arbitrarily small length scales become amplified, and no stable pattern is formed. We thus only search for Turing I instabilities in the following. See Figure 1A for a visualization of this phenomenon and the sections below for mathematical details.

Therefore, to find Turing instabilities we first need to identify the stable steady states of a given system, and subsequently study their dispersion relation. Figure 2C summarizes the computational procedure.

For each candidate network we perform this procedure for a wide range of kinetic and diffusion parameters. We therefore have to specify the intracellular parameters determining the regulatory functions *f*_*i*_, as well as the diffusion constants. The former consist of the parameters k (dissociation rates), V (scaling factors), b (basal production rates) and the degradation rate *µ* (see Equations in Figure 2B), which we vary across biologically relevant values between 0.1 and 100 (0.01-1 for *µ*). The extracellular/diffusion parameters are varied over a range between 10^−3^ and 10^3^. Due to computational cost, we vary the V and b parameters only for 2-node networks, but fix them to 100 and 0.1, respectively, for 3-node networks. In this way we were still able to screen 3 × 10^11^ network-parameter combinations for TP formation, which we believe is the largest study of its kind to date.

### Definition of networks and ODEs

#### Networks

To generate all possible *n*-node networks we first compute all possible (*n* × *n*)–matrices with elements 0, 1 and −1, where a 0 (1/ −1) represents the absence (presence) of an activating or inhibiting interaction/edge. The number of matrices is reduced by considering only connected networks and accounting for symmetries. We also remove matrices that correspond to networks including nodes without any incoming or outgoing edges. Each remaining matrix *M* serves as an adjacency matrix for a network, where the element *M*_*ij*_ being 1 (−1) represents a positive (negative) edge from node *i* to node *j*.

#### System of ODEs

For each adjacency matrix we then construct the corresponding set of ODEs. Each non-zero entry in the adjacency matrix corresponds to a Hill-type term which are combined in either a non-competitive or competitive case. If 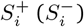 denotes the set of positive (negative) edges ending in node *i*, then in the non-competitive case, the different regulators act (and saturate) independently of each other, and we have,

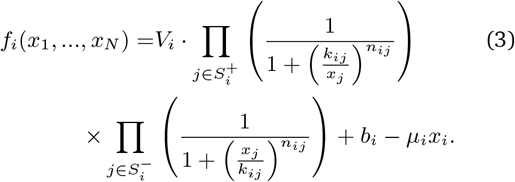

Here, *V*_*i*_ is the maximal induced production rate, *b*_*i*_ the basal production rate, *µ*_*i*_ the degradation rate, *k*_*ij*_ the concentration value at which the regulation of the *i*th species by the *j*th species is half its maximal value, and *n*_*ij*_ is the corresponding Hill coefficient determining the steepness of the response.

In the competitive case the different regulators compete for the binding site which leads to an additive combination of terms:

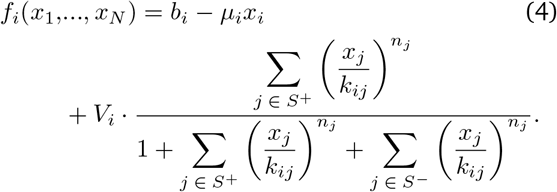

### Numerical analysis

#### Steady state estimation

We generated a customized Matlab (R2016a) script to find steady states numerically. For a given system the parameters are chosen from a logarithmic grid and for each set of parameters the system of ODEs is solved numerically until time *t* = 1000 using the Matlab ODE solver ode15s. Whenever the algorithm encounters numerical problems, the (slower but more robust) algorithm ode23s is invoked.

The resulting trajectory is checked to have converged to a steady state. Next, we solve the system again for 3*n* initial conditions (Geest, 2016), where *n* is the number of species. The endpoints of each resulting trajectory are clustered using *k*-means clustering assuming that the steady states are given by the centroids of the resulting clusters. Using the cluster centroids as an initial condition, the system of ODEs is solved again to verify that the system has converged sufficiently. For steady states that are approached via damped oscillations, we do not solve the ODEs numerically but instead search for a fixed point of the ODEs directly using the Matlab function fsolve.

#### Stability Analysis

The stability of a steady state **x**\* of Equation (1) is assessed by a linear stability analysis. We add a small perturbation around **x**\*

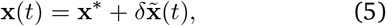

with a small constant *δ* ∈ ℝ, and then linearise Equation (1), to obtain

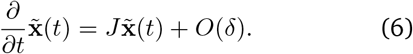

The Jacobian, *J*, is defined as

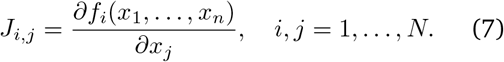

Equation (6) constitutes a linear dynamical system with steady state 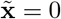. This steady state is asymptotically stable if and only if the real parts of all eigenvalues of the matrix *J* are negative. This in turn means that **x**\* is a locally asymptotically stable steady state, in the sense that there exists a neighborhood around **x**\* such that any solution of the ODEs starting from this neighborhood asymptotically converges to **x**\*.

Accordingly, it is sufficient to compute the eigenvalues of the Jacobian defined in Equation (7) to assess the local stability of a steady state of Equation (1).

To assess if such a stable steady state can exhibit a diffusion-driven instability, we need to analyze Equation (2). Similarly to the case without diffusion, we perturb the system around **x**\* to perform a linear stability analysis of this steady state, but this time with a harmonic wave with wave-length *q*:

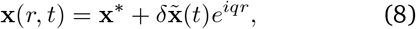

with a small constant *δ* ∈ ℝ. Inserting this into Equation (2) and expanding to first order in *δ*, one obtains a linear dynamical system similar to the one in (6), but with a modified Jacobian 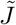 given by:

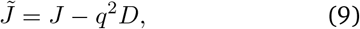

where *D* = diag(*D*_1_, …, *D*_*n*_) is a diagonal matrix with the diffusion constants *D*_*i*_ on the diagonal. For differential diffusion of the molecular species, the Jacobian in Equation (8) can have eigenvalues with positive real part for finite wave vector *q*; when this occurs the stable steady state of the system in Equation (1) becomes unstable with diffusion, and we speak of an diffusion-driven Turing instability. Depending on the behavior of the largest eigenvalue of 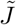 for large *q* we distinguish different types of Turing instabilities, see Figure 4D.

#### Workflow

We implement the estimation of stable steady states of the non-spatial system and identification of Turing instabilities into an automated workflow. Overall, the computational analysis consists of the steps:

i. Define a network and its governing ODE equations.
ii. Define a grid that covers the full parameter space of interest.
iii. For each set of parameters:

a. Perform numerical simulation of ODE system for different initial conditions
b. Cluster endpoints using k-means clustering to find set of steady states.
c. Confirm steady state guesses by repeated simulation from each cluster center.
iv. Perform stability analysis for all steady state and parameter combinations:

a. Calculate Jacobian matrix (without diffusion) and test if all real parts of the eigenvalues are negative to verify stability.
b. Expand Jacobian matrix by diffusion term and calculate eigenvalues for a set of diffusion values and wavenumbers q.
c. Classify possible Turing instabilities as Type Ia, Ib, or IIa or IIb (see Figure 2C).

### Software package

We provide code for an automated steady state estimation an stability analysis together with documentation in Supplementary File S4. The code can analyze systems with arbitrary regulatory functions and node numbers.

